# An economic, square-shaped flat-field illumination module for TIRF-based super-resolution microscopy

**DOI:** 10.1101/2021.11.05.467489

**Authors:** Jeff Y.L. Lam, Yunzhao Wu, Eleni Dimou, Ziwei Zhang, Matthew R. Cheetham, Markus Körbel, Zengjie Xia, David Klenerman, John S.H. Danial

## Abstract

Super-resolution (SR) microscopy allows complex biological assemblies to be observed with remarkable resolution. However, the presence of uneven Gaussian-shaped illumination hinders its use in quantitative imaging or high-throughput assays. Methods developed to circumvent this problem are often expensive, hard-to-implement, or not applicable to total internal reflection fluorescence (TIRF) imaging. We herein demonstrate a cost-effective method to overcome these challenges using a small square-core multimodal optical fibre as the coupler. We characterise our method with synthetic, recombinant and cellular systems imaged under TIRF and highly inclined and laminated optical sheet (HILO) illuminations to demonstrate its ability to produce highly uniform images under all conditions.

## 1. Introduction

SR microscopy has led to fundamental breakthroughs in biology by allowing cellular structures smaller than the diffraction limit of light to be readily observed [1–3]. Single-molecule localisation microscopy (SMLM) techniques, such as (direct) stochastic optical reconstruction microscopy ((d)STORM) [4], photoactivated localisation microscopy (PALM) [5] and DNA points accumulation for imaging in nanoscale topography (DNA-PAINT) [6], achieve SR by switching the emission of individual fluorophores one-at-a-time and recording the position of each fluorophore down to a single-digit nanometre precision through the fitting of a point spread function. By virtue of its high resolving power and versatility, SMLM has been a highly popular tool employed worldwide.

To fully utilize the camera chip for imaging large samples by SMLM techniques, a full field-of-view (FOV) needs to be illuminated. As such, illumination homogeneity crucially affects the quality of a super-resolved image [7–9]. In (d)STORM and DNA-PAINT, the on-rate of fluorophores depends heavily on the intensity of the excitation source [7,9]. Hence, uneven illumination results in a non-uniform distribution of emitted photons across the FOV and unwanted imaging artefacts. Poorly illuminated regions give rise to unclear structures, whilst strongly illuminated regions result in imprecise localizations that hamper quantitative analysis and molecular counting [7,10]. In conventional SMLM setups, the excitation source is coupled into the objective by free-space optics leading to a non-homogenous Gaussian-shaped illumination profile that induces the above problems. Recent attempts were made to reshape Gaussian-shaped beams into uniform flat-field illumination sources using microlens arrays [7], diffractive beam shaping devices [10], refractive beam shapers [9], and optical waveguides [11]. Unfortunately, the application of these methods is hindered by either high cost or complicated optical setup.

Multi-mode fibres (MMFs) provide alternative means to obtaining flat-field illumination. As MMFs allow the transmission of multiple spatial modes, they produce even illumination at a high coupling efficiency [8]. MMFs circumvent the drawbacks of using free space based flat-field illumination including the high cost and requirement for fine alignment of laser lines at the sample plane. It was earlier reported that a small circular core (50 μm and 105 μm) MMF could deliver a flat-field illumination suitable for TIRF imaging but at the expense of producing a non-square FOV as well as non-homogenous beam profile due to poor mode scrambling [12]. More recently, a large square-core (150 μm x 150 μm) MMF was used to deliver a square flat-field illumination pattern for epi-based experiments but not applicable under TIRF imaging due to the large size of the fibre core diameter [13]. Imaging under TIRF can potentially give better signal-to-noise ratio as only the fluorophores bound on the surface within the evanescent field will selectively be excited. This is especially important when there is an elevated background fluorescence, such as in DNA-PAINT imaging. Yet, none of the developed solutions with MMF can deliver a square flat-field illumination that is suitable for surface-based SR imaging.

In this study, we demonstrate an economic flat-field illumination for TIRF-based SR microscopy by utilising a small square-core (70 μm x 70 μm) MMF in the combined excitation path. There are two advantages to using a square-core fibre:

1. it enables the full utilisation of the square and rectangular camera chips;
2. it delivers improved mode scrambling compared to circular-core fibres [13]. This is fundamentally important for minimizing the localization precision, improving stoichiometry quantification across the FOV, as well as for the faithful representation of the underlying biological samples.

We systematically characterise the performance of our MMF with various samples, including fluorescent microspheres, CellMask™ Orange Plasma membrane stained-T cells, supported lipid bilayers, synthetic DNA origamis, cellular microtubules, and recombinant protein aggregates illuminated under HILO and TIRF. Our versatile and cost-efficient system can be easily assembled, aligned, and maintained to provide a square-shaped flat-field illumination.

## 2. Results

To achieve a square-shaped flat-field illumination profile, we introduced a small square-core (70 μm x 70 μm) MMF in the combined excitation path of our home-built TIRF microscope (fig. 1. a, see methods). Briefly, the Gaussian-shaped output of four laser lines was coupled into the fibre using an aspheric lens at high coupling efficiencies (81% for 405 nm, 86% for 488 nm, 93% for 561 nm and 82% for 638 nm). The output of the fibre was then collimated and focused onto the back focal plane of the objective using a combination of an adjustable-focus fibre collimator and an achromatic doublet lens to produce a square-shaped flat-field illumination profile at the sample plane (fig. 1. b). Autofluorescence generated by the fibre was removed using a quad-band excitation filter and speckles caused by interference were eliminated using a cylindrical vibration motor attached onto the MMF with a 3D-printed mount (fig. 1. a, fig. S1. and S2., see methods).

**Fig. 1.**
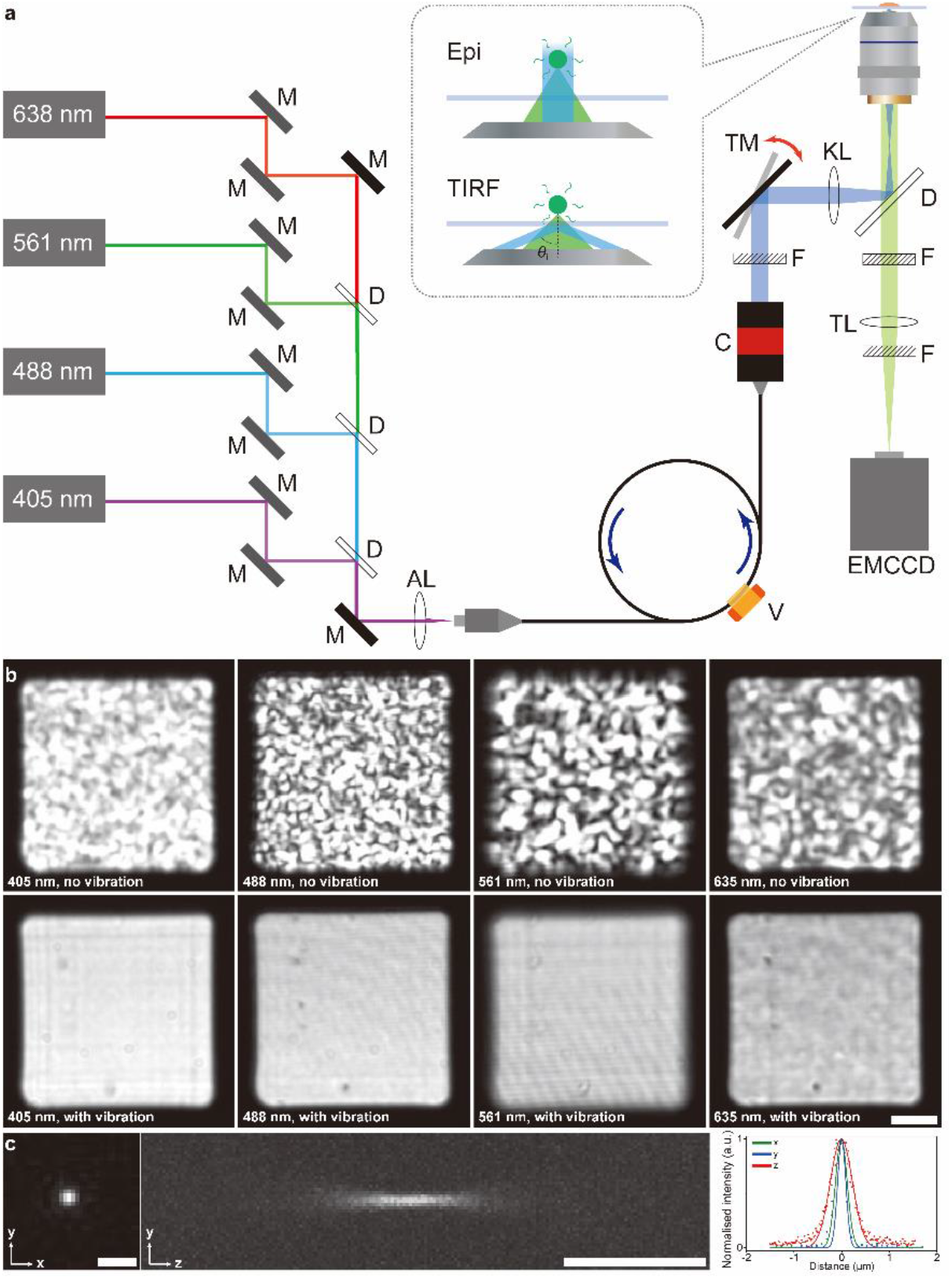
Schematic diagram of our home-built TIRF microscope with flat-field illumination and its demonstration as well as resolution characterisation. (a) Schematic diagram of the setup. Solid black filled rectangles M represent dielectric mirrors for aligning; C represents the fibre collimator; V represents the vibration motor; TM shows the dielectric mirror for adjusting the angle of incident beam *θ*; black unfilled rectangles D represent dichroic mirrors; striped rectangles F represent filters; ellipses AL, KL and TL represent aspheric lens, achromatic doublet lens, and tube lens, respectively. The single arrow indicates the direction of the combined laser beam while the double arrow indicates the alternation of TM. (b) Demonstration of the illumination profile after coupling into the square-core optical fibre without (top) and with (bottom) fibre agitation (i.e. vibration) for speckle reduction as imaged on a CMOS camera (DCC1545M-GL, Thorlabs) after collimation for all four different lasers. Scale bar, 10 μm. (c) Point spread function (PSF) of our home-built TIRF microscope with flat-field illumination, measured by 100 nm microspheres under 488 nm excitation. Representative cross-sections of the PSF of the fluorescent microspheres are shown in the *x-y* plane (left) and in the *y-z* plane (middle). Intensity plots along the three axes (right) show the full width at half maximum (FWHM) are 293.4 nm, 272.1 nm and 544.2 nm along the *x, y*, and *z* axes, respectively. Scale bars, 1 μm.

We first assessed the resolution of our flat-field illumination module by calculating the full-width at half-maximum (FWHM) of the effective point spread function (PSF) obtained from 100 nm microspheres under 488 nm excitation. The measured lateral resolutions of our system were 294 ± 22 nm in the *x*-direction and 286 ± 20 nm in the *y*-direction, which are comparable to the resolutions of most conventional TIRF microscopes [14]. The axial resolution of our system was 545 ± 15 nm (fig. 1. c).

In our inverted fluorescence microscope system, the angle of the incident laser beam (*θ* in fig. 1. a) can be varied easily with a mirror (TM in fig. 1. a) at the back of the microscope. This allows us to switch between epi-illumination, HILO and TIRF. With the application of a small core optical fibre, the full laser beam can be totally reflected without any partial refraction that could otherwise cause unintended excitation of fluorophores in the background under TIRF-based imaging.

We then evaluated the homogeneity of the emission profile by imaging a supported lipid bilayer, a smooth surface consisting of two layers of lipid molecules that exhibits lateral fluidity, formed on a glass coverslip. To accomplish this, we doped 1% fluorescently labelled Oregon Green-1,2-dihexadecanoyl-*sn*-glycero-3-phosphoethanolamine (OG-DHPE) into 1-palmitoyl-2-oleoyl-*sn*-glycero-3-phosphocholine (POPC). This gives rise to a uniformly fluorescent surface and therefore allows us to observe the emission profile of the microscope under Gaussian and square-shaped flat-field illuminations (fig. 2. a and b, respectively). Examining the acquired images, we uncovered a two-fold intensity difference between the centre and peripheries of the FOV under conventional Gaussian-shaped illumination (fig. 2. a (bottom)), while observing a highly uniform emission profile across the entire FOV under flat-field illumination (fig. 2. b (bottom)).

**Fig. 2.**
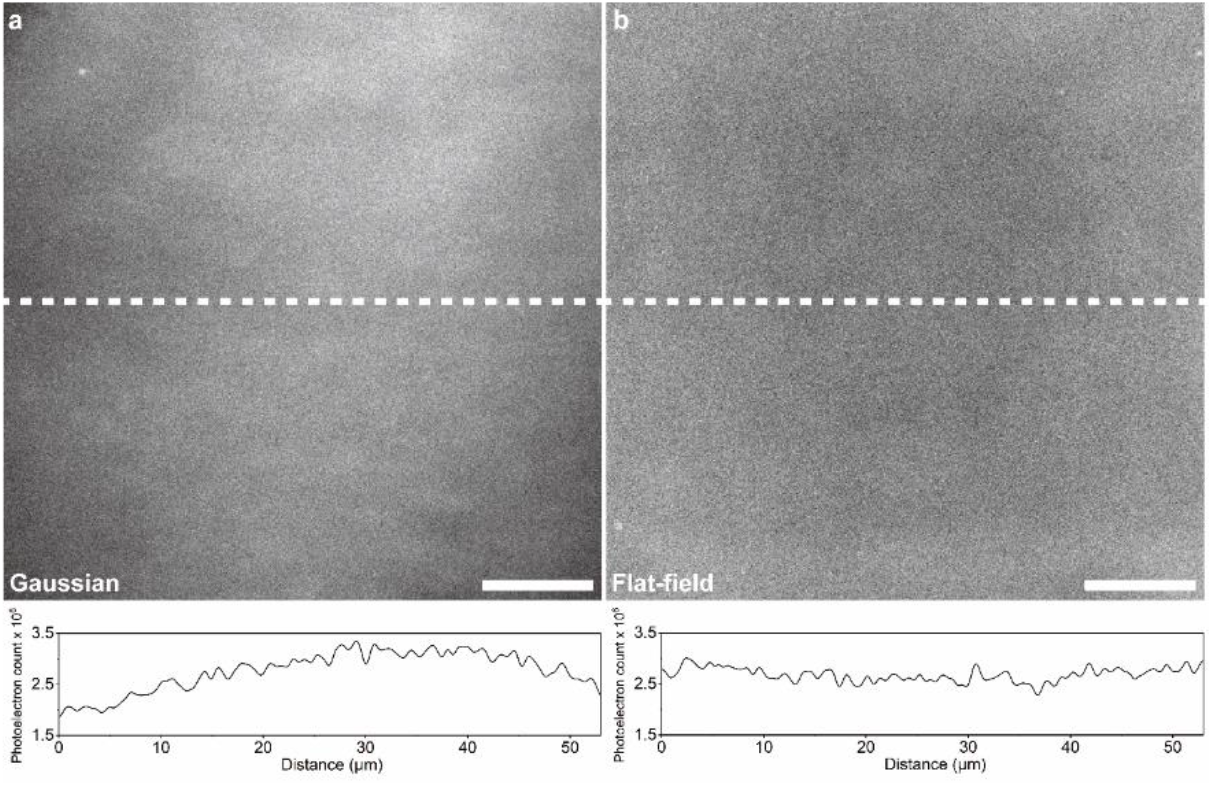
TIRF images of the full FOV (top) and line plots of fluorescence intensity along the principal axis (white, bottom) obtained by supported POPC lipid bilayer with 1% OG-DHPE under (a) conventional Gaussian and (b) flat-field illumination profiles. Although the ‘Gaussian’ profile represents the profile of a free-launched beam, the profile deviates from a perfect Gaussian shape as is common for some lasers. Scale bars, 10 μm.

We first performed a comparative assessment of the suitability of our MMF for TIRF by imaging surface-adhered CellMask™ Orange Plasma membrane stained-T cells using a 50 μm circular core MMF (solution proposed in [12], fig. 4. a and b), 70 μm square core MMF (solution proposed here, fig. 4. c and d), and 200 μm square core MMF (solution proposed in [13], fig. 4. e and f). The 50 μm circular core, and 70 μm square core MMFs show higher quality TIRF imaging compared to the 200 μm square core MMF which shows the appearance of membrane contours (fig. 4. f) out of focus. Furthermore, the 50 μm circular core fibre shows an uneven illumination pattern (fig. 4. a) compared with the 70 μm fibre (fig. 4. c), which is presumably due to poor mode scrambling. Our theoretical calculations (see supplementary information) show that our microscope objective has a 310 μm TIRF annulus and that the beam size at the back focal aperture is 262.5 μm. This confirms that the beam in its entirety contributes to production of an evanescent field for TIRF imaging.

We then assessed the suitability of our system for SR imaging using DNA-PAINT. Successful DNA-PAINT imaging requires the sample of interest to be excited under TIRF to suppress the background of fluorescently labelled imaging strands and illuminate the surface-bound species alone. To demonstrate the applicability of our fibre-based solution to TIRF-based DNA-PAINT imaging, we imaged surface-immobilised nanorulers (three-spot DNA origami nanostructures with 40 nm distance between each spot, fig. 3. a) under epi- and TIRF illumination (fig. 5. a and b). Poor signal-to-noise ratio due to high background fluorescence was observed under epi-illumination (fig. 5. a); on the other hand, improved signal-to-noise ratio was obtained with TIRF as the fluorescence intensity of the fluorophore bound on the surface was enhanced whilst that in the bulk outside the evanescent field was dramatically reduced (fig. 5. b). This again demonstrates the ability of our fibre-based flat-field illumination module for TIRF-based imaging. Furthermore, we could resolve the nanorulers with fiducial markers across the entire FOV (fig. 5. c) at a mean localisation precision of 8.7 nm (fig. 5. d). The image resolution was calculated to be 35.5 nm by Fourier ring correlation (FRC) (fig. 5. e). This demonstrates the ability of our setup to resolve nanoscopic structures with exceptional uniformity under TIRF illumination.

**Fig. 3.**
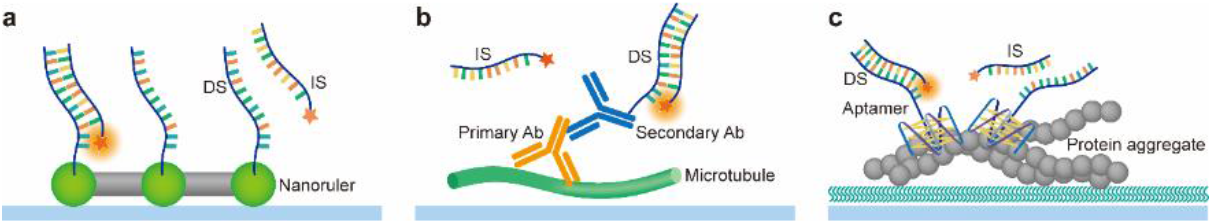
Schematic diagrams for DNA-PAINT with (a) DNA origami nanostructures on nanorulers, (b) fixed cellular microtubules immunostained by DNA-antibody conjugates and (c) aptamer (AD-PAINT) targeting protein aggregates. DS – docking strand; IS – imaging strand; Ab – antibody.

**Fig. 4.**
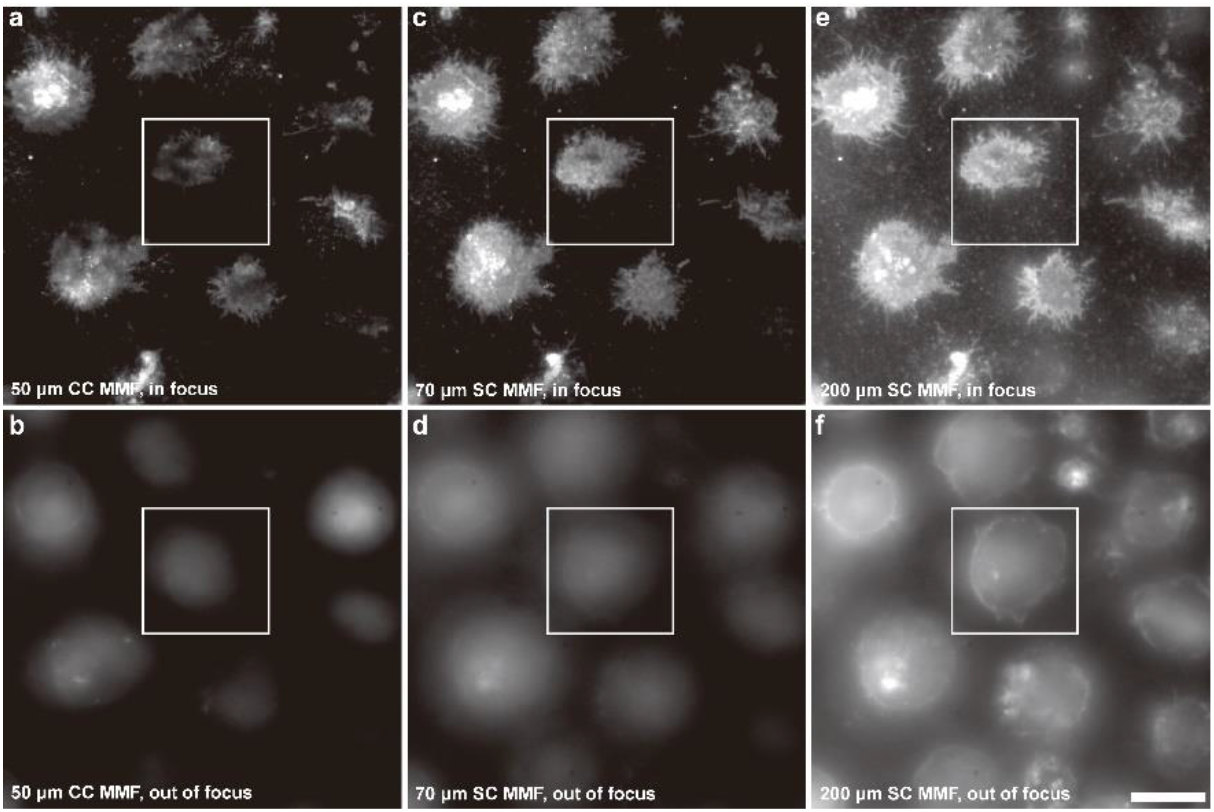
Diffraction-limited images of surface-adhered CellMask™ Orange Plasma membrane stained-T cells illuminated using a 561 nm laser line coupled into 50 μm circular core (CC) MMF (solution proposed in [12], a and b), 70 μm square core (SC) MMF (solution proposed here, c and d), and 200 μm square core MMF (solution proposed in [13], e and f). Images were acquired at the focal plane (top) and 4 μm above the focal plane (bottom). Scale bar, 10 μm.

**Fig. 5.**
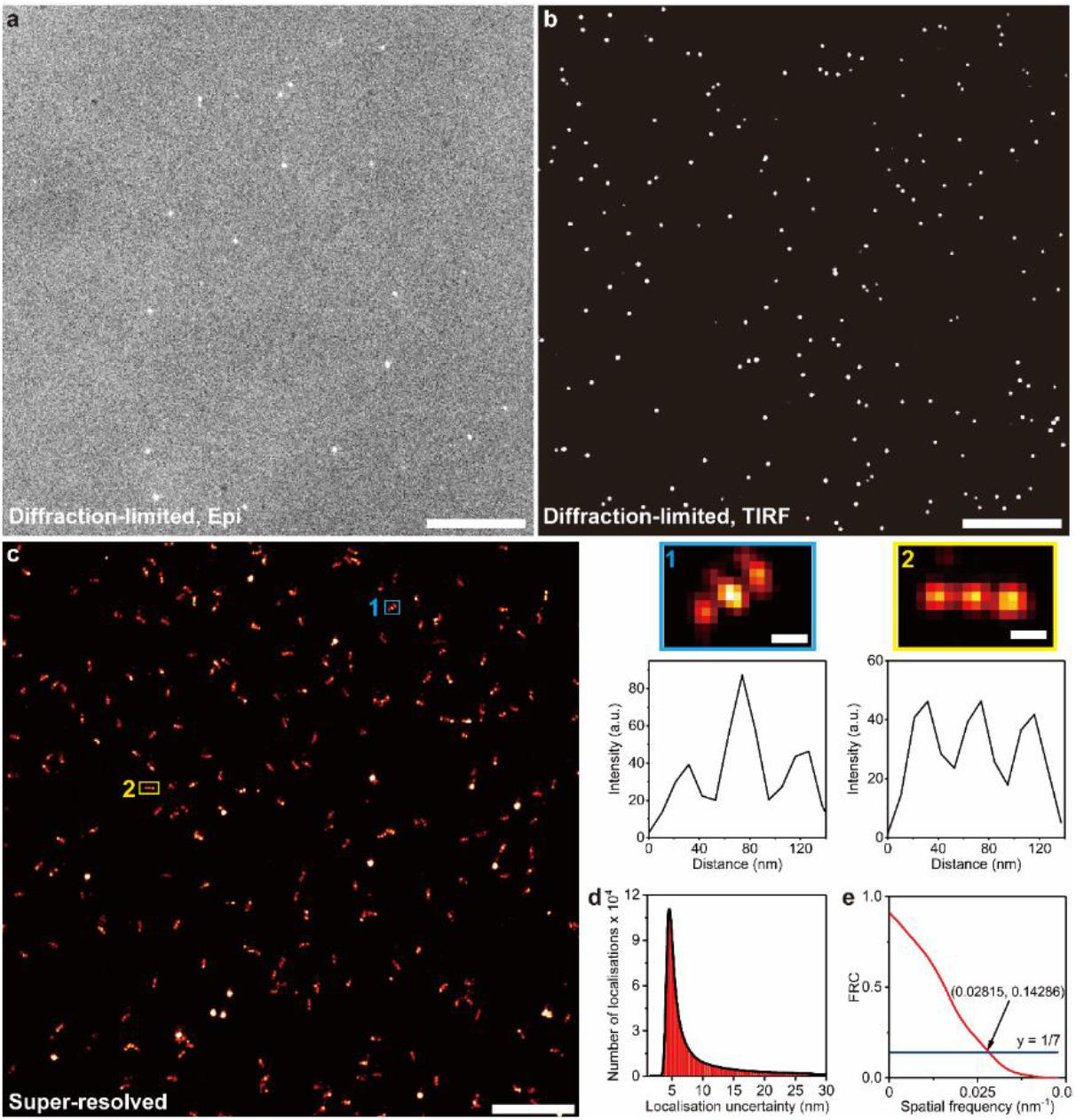
Diffraction-limited and super-resolved images of the nanorulers and their analyses. Diffraction-limited images of nanorulers under (a) epi- and (b) TIRF illumination. Both images were taken at the same plane with the same exposure settings and adjusted to the same level of brightness and contrast. A clear high background fluorescence (approximately 6-fold) is observed (a) under epi-illumination compared to (b) TIRF illumination. Scale bars 10 μm (a and b). (c) Super-resolved image of the three-spot DNA origami nanostructures with 40 nm distance between each spot with imaging strand attached to cy3B, with a laser power of 111 mW before the objective. The image was reconstructed and drift-corrected from 10000 frames with an exposure time of 100 ms. Scale bar 1 μm. (1) and (2) Close-up images of the corresponding regions in (c) (top, scale bar 50 nm) and their corresponding intensity vs distance profile (bottom). (d) The number of localisations vs. the localisation uncertainty shows that we can achieve DNA-PAINT imaging with the nanorulers at a precision of 8.7 nm. (e) The Fourier ring correlation (FRC) of the DNA-PAINT imaging of the nanoruler. This shows that the resolution obtained was 35.5 nm at which the curve drops below 1/7.

With the success of applying our fibre-based solution to imaging synthetic surface-based structures under TIRF illumination, we then used our system for visualising cellular microtubules (fig. 3. b) under HILO illumination to interrogate the entire cell volume. The microtubules were immunostained using antibodies either conjugated with dyes for dSTORM, or docking strands for DNA-PAINT (see methods). Using our fibre-based, square-shaped flat-field module, a uniform localisation density was maintained from the centres to the peripheries of the FOV in both DNA-PAINT imaging (fig. 6. a) and dSTORM imaging (fig. 6. b) under HILO. The images confirm the suitability of our solution for resolving nanoscopic cellular structures precisely under HILO illumination and uniformly across a wide FOV.

**Fig. 6.**
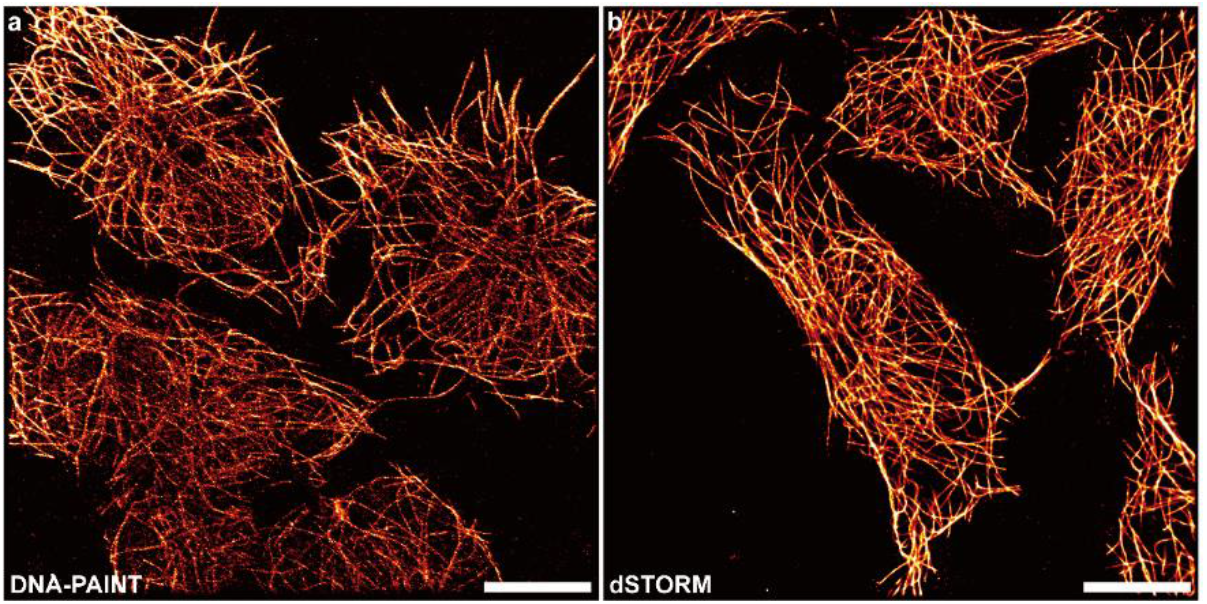
Super-resolved, full FOV image of the microtubules in fixed and permeabilised HEK293T cells for (a) DNA-PAINT imaging and (b) dSTORM under HILO illumination in our fibre-based, square-shaped flat-field setup. Scale bar 10 μm.

Having demonstrated the applicability of our fibre-based system to imaging synthetic and cellular samples, we applied it for imaging recombinantly prepared protein aggregates (figure 3c) using aptamer DNA-PAINT (AD-PAINT). AD-PAINT is a recently developed technique for examining surface-captured protein complexes below the diffraction limit [15]. Aptamers are single-stranded oligonucleotides that can fold into G-quadruplexes, a secondary nucleic acid structure formed among guanine bases by Hoogsteen type hydrogen bond. Some aptamers have high affinity and specificity for different protein oligomers and fibrils, thus acting as tiny molecular probes [16]. By introducing a DNA docking strand on an aptamer, which is complementary to a fluorescently labelled imaging strand, it is possible to perform DNA-PAINT on surface-captured protein aggregates. Here, we used an aptamer that recognises fibrils of the intrinsically disordered protein α-synuclein which is implicated in Parkinson’s disease [15–17]. We first imaged the fibrils at the diffraction limit using thioflavin-T, a dye that interacts with the hydrophobic regions of protein aggregates, to ensure their presence (fig. 7. a) then super-resolved them using AD-PAINT (fig. 7. b) all under TIRF illumination. As with the other samples, we observe uniform imaging of the fibrils across the entire FOV demonstrating the suitability of our solution for high-throughput light-based assays.

**Fig. 7.**
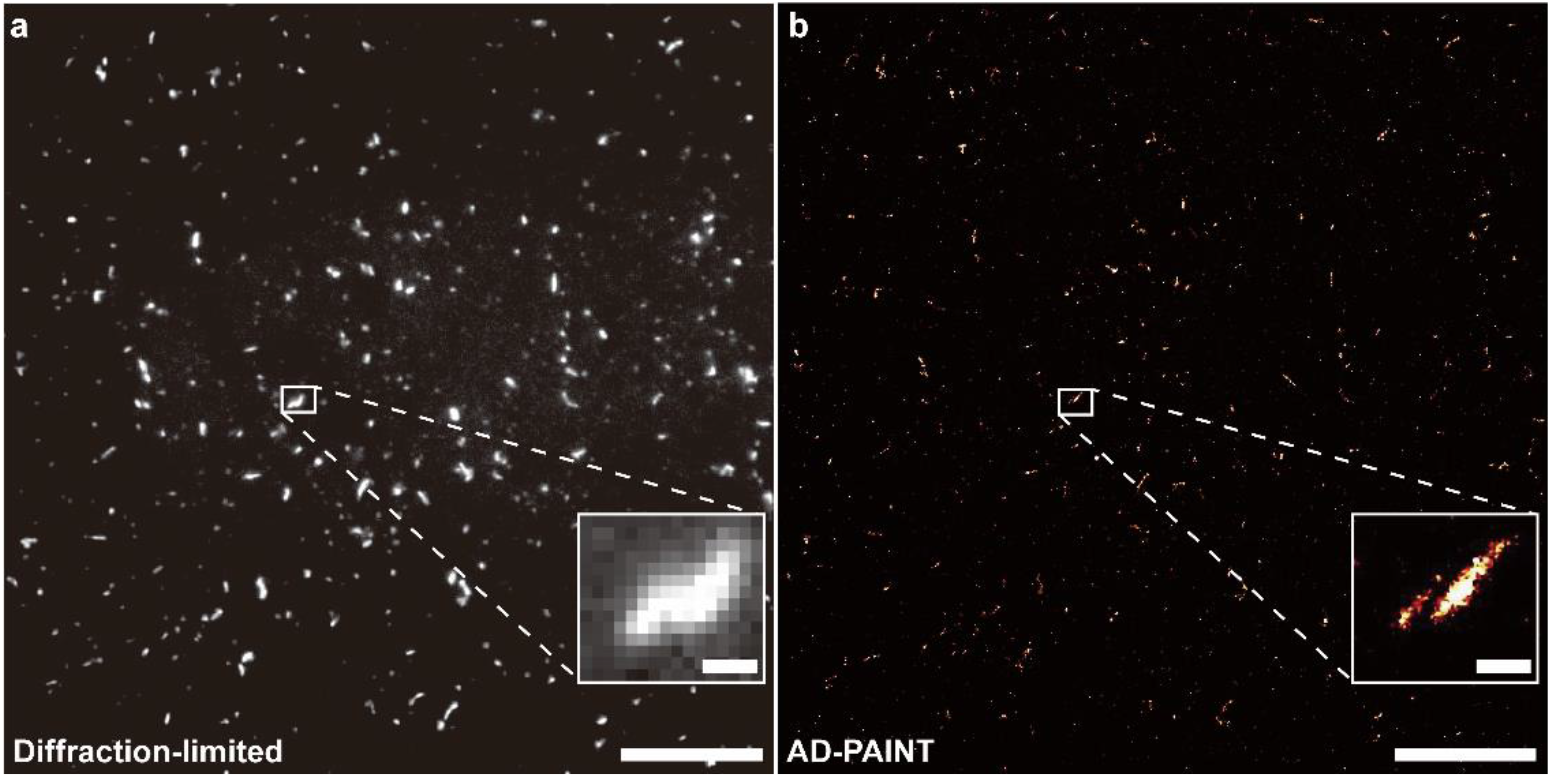
(a) Diffraction-limited and (b) super-resolved image of the α-synuclein fibrils with thioflavin-T and aptamer-docking strand DNA respectively. The scale bars for (a) and (b) are 10 μm while that of the close-up in the (a) diffraction limited images and (b) super-resolved images are 500 nm.

## 3. Discussion

Currently available options to generate a square flat-field illumination are dependent on the use of expensive beam shapers which are not compatible with commercially-available, fibre-based laser engines or large core, square-core fibres which are only compatible with epi- / HILO illumination, but not TIRF. Our small square-core MMF is:

1. compatible with fibre-based laser engines (which are widely used in commercially-available or home-built setups [18]),
2. economic (1000 USD compared to 5000 USD for a beam shaper [10]),
3. compatible with TIRF illumination (owing to the small size of its core compared to widely-available square-core MMFs [13]),
4. optimized for square and rectangular large camera chips (owing to the square core and its ability to illuminate large FOVs),
5. compatible with highly-homogenous illumination (owing to improved mode scrambling compared to circular-core MMFs [13]).

Our solution is novel; not only adding important value and innovation to the field, but setting a standard to high-quality imaging, at low-cost, and with very easy implementation. The benefits that this solution delivers are disproportionally more compared to existing solutions, and we expect its quick adoption across academia and industry.

## Funding

J.Y.L.L. is supported by the Croucher Foundation Limited (Hong Kong). E.D is funded by a Deutsche Forschungsgemeinschaft (DFG) Research Fellowship (426806622). M.K. is funded by the Royal Society (RP/EA/180002). J.S.H.D is funded by a postdoctoral fellowship from EISAI and the UK Dementia Research Institute (DRI) Limited, pilot grant from the UK DRI Ltd and a research associateship from King’s College, University of Cambridge. D.K. is funded by a European Research Council (ERC) advanced grant (669237), the UK DRI Ltd funded by the UK Medical Research Council (MRC), and by the Royal Society (UK).

## Acknowledgments

We would like to thank Jonas Ries (EMBL) for useful preliminary discussions. We would also like to thank Simon J. Davis and Edward Jenkins (University of Oxford) for providing the J8 LFA-1 Jurkat T-cell line. J.S.H.D conceived and designed the study. J.Y.L.L, Y.W and Z.X constructed the microscopy setups. J.Y.L.L, Y.W, E.D, Z.Z, M.R.C and M.K prepared all samples. J.Y.L.L and Y.W performed all analysis. J.Y.L.L wrote the manuscript with input from all authors. D.K supervised the project.

## Disclosures

The authors declare no competing interests.

## Supplemental document

See Supplemental document for supporting content.

## Notes

### Competing Interest Statement

The authors have declared no competing interest.

