## Supplementary Information for "An economic, square-shaped flat-field illumination module for TIRF-based super-resolution microscopy"

### 1. Methods

#### *Home-built TIRF microscope with flat-field illumination setup*

Four lasers operating at 405 nm (LDM-405-350-C, Lasertack GmbH), 488 nm (Toptica iBeam smart, Toptica), 561 nm (Cobolt Jive, HÜBNER GmbH & Co KG) and 638 nm (Cobolt 06-MLD-638, HÜBNER GmbH & Co KG) were coupled to the optical axis of a 1.49 N.A. 100x CFI Apo TIRF objective (MRD01991, Nikon) mounted on an inverted Ti-E Eclipse microscope (Nikon, Japan). The laser powers were controlled by their corresponding software or attenuated by neutral density filters. The laser beams then passed through the aligning mirrors and were combined by their corresponding dichroic mirror (for 405 nm: FF458-Di02-25x36, Semrock; for 488 nm: FF552-Di02-25x36, Semrock; for 561 nm: FF605-Di02-25x36, Semrock) before being focused by an aspheric lens (C220TMD-A, Thorlabs) to the square-core optical fibre (05806-1 Rev. A, CeramOptec). The combined laser beam coming out from the optical fibre was then collimated (C40FC-A, Thorlabs) and cleaned up by a quad-band excitation filter (FF01-390/482/563/640-25x36) before passing through the back port of the microscope and reaching the achromatic doublet lens (AC254-125-A-ML, Thorlabs). Speckles from the fibre were removed using a vibration motor mounted on a custom 3D printed mount (Figures S1 and S2). The excitation beam was then reflected by a quad-band dichroic beam splitter (Di01-R405/488/561/635-25x36, Semrock) or a penta-band dichroic beam splitter (R405/488/561/635/800-T1-25x36, Semrock) and focused on the sample by the objective. Fluorescence from the sample was collected by the objective and passed through the quad-band or penta-band dichroic beam splitter. The fluorescence was then cleaned up with a quad-band emission filter (FF01-446/523/600/677-25x36, Semrock) and their corresponding appropriate filters (for both 405 and 488 nm induced fluorescence: BLP01-488R-25x36, Semrock and FF01-520/44-25x36, Semrock; for 561 nm induced fluorescence: LP02-568RS-25x36, Semrock and FF01-587/35-25x36, Semrock; for 638 nm induced fluorescence BLP01-635R-25x36, Semrock) mounted on a high-speed filter wheel (HF110A, Prior Scientific) before being recorded on an EMCCD camera (Evolve 512, Photometrics) operating in frame transfer mode (EM Gain of 3.1 electrons/ADU and 250 ADU/photon). Each pixel corresponds to a length of 105.4 nm, 106.3 nm or 101.2 nm on the recorded image. The microscope was also fitted with a perfect focus system (PFS) which auto-corrects the *z*-stage drift during a prolonged period of imaging. To remove the stray infrared laser beam, a short-pass filter (FESH0750, Thorlabs) was mounted on the entrance port of the EMCCD camera.

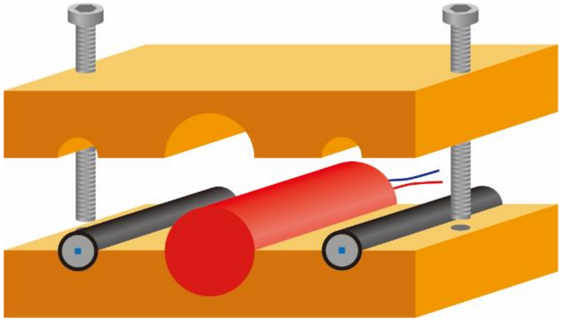

Figure S1. Schematic diagram of the vibrator in the home-built TIRF microscope with flat-field illumination.

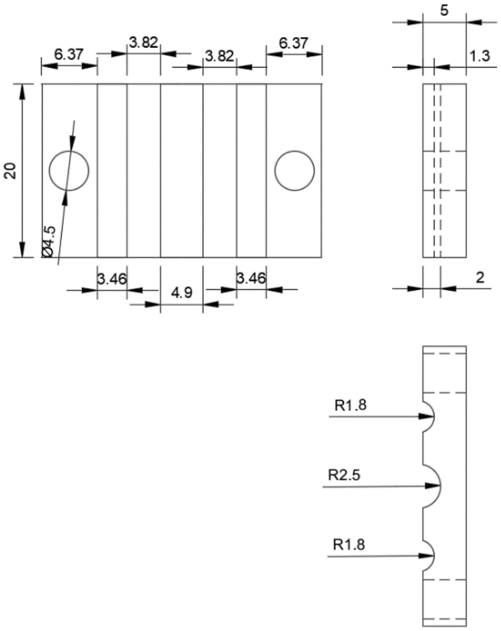


Figure S2. 2D sketch of the vibrator mount in the home-built TIRF microscope with flat-field illumination. All measurements are in millimetres. R – radius.

#### *Home-built TIRF microscope with Gaussian illumination setup*

A laser operating at 488 nm (Toptica iBeam smart, Toptica) was coupled to the optical axis of a 1.49 N.A. 100x CFI Apo TIRF objective (MRD01991, Nikon) mounted on an inverted Ti-E Eclipse microscope (Nikon, Japan). The laser power was controlled by TOPAS iBeam smart 1.4.0.115 or attenuated by neutral density filters. The laser beam then passed through a quarter-wave plate for circular polarisation and was cleaned up by an excitation filter (LL01-488-25x36, Semrock). The laser beam was then expanded by a beam expander before reaching a dichroic mirror (FF552-Di02-25x36, Semrock). Next, the reflected laser beam was focused by an achromatic doublet lens (AC254-400-A, Thorlabs) and reflected by a quad-band dichroic beam splitter (Di01-R405/488/561/635-25x36, Semrock). The objective then focused the reflected excitation beam on the sample. Fluorescence was collected by the objective and passed through the quad-band dichroic beam splitter. It was then cleaned up by an emission filter (BLP01-488R-25x36, Semrock) before being recorded on an EMCCD camera (Evolve 512, Photometrics) operating in frame transfer mode (EM Gain of 3.1 electrons/ADU and 250 ADU/photon). Each pixel corresponds to a length of 101.6 nm on the recorded image. The microscope was also fitted with a perfect focus system (PFS) which auto-corrects the *z*-stage drift during a prolonged period of imaging. To remove the stray infrared laser beam, a short-pass filter (FESH0750, Thorlabs) was mounted on the entrance port of the EMCCD camera.

#### *Imaging conditions*

Slides were fixed on a microscope stage and coupled to an objective using refractive index-matched low-autofluorescence immersion oil (refractive index $n$ = 1.518, Olympus, UK). Images were taken in a grid using an automation script (μManager v1.4.22). Exposure times were set at 50 ms for diffraction-limited images for α-synuclein fibrils with thioflavin-T, 200 ms for diffraction-limited images for CellMask$^{TM}$ Orange Plasma membrane stained-T cells, 33 ms for dSTORM, 100 ms for DNA-PAINT for DNA origami nanoruler (40 nm, cy3B, (40Y), immobilised high-resolution DNA-PAINT nanoruler, GATTAquant GmbH, Cat. No. 3030) and 150 ms for DNA-PAINT of microtubules in cells and AD-PAINT. For DNA-PAINT imaging of the nanoruler, 11000 frames were acquired; for cellular dSTORM and DNA-PAINT imaging, 45000 and 60000 frames were acquired, respectively; for AD-PAINT imaging, 10000 frames were recorded; for diffraction-limited images for α-synuclein fibrils with thioflavin-T and CellMask$^{TM}$ Orange Plasma membrane stained-T cells, 50 and 100 frames were acquired respectively.

#### *Resolution characterisation of home-built TIRF microscope with flat-field illumination setup with fluorescent microspheres*

TetraSpeck™ microspheres, 0.1 μm, fluorescent blue/green/orange/dark red (ThermoFisher, T7279) were embedded in 2% agarose gel (Merck, Cat. No. A9414) on a glass coverslip in a ratio of 1:500. The sample was then excited by a 488 nm laser. The raw images were sliced and FWHMs were calculated to determine the optical resolution of our setup. The final calculated FWHMs were obtained by averaging 10 beads of different FOVs.

#### *Imaging of beam profile*

Three 15 cm extension tubes (SM1E60, Thorlabs) were connected and installed in front of the collimator in our home-built TIRF microscope with flat-field illumination setup. The beam profiles from different laser lines were recorded with a CMOS camera (DCC1545M-GL, Thorlabs) and the images were acquired with ThorCam software.

#### *Preparation of lipid vesicle suspension*

To 40 μL of 25 mg/mL 1-palmitoyl-2-oleoyl-sn-glycero-3-phosphocholine (POPC, Avanti, Cat. No. 850457C-200mg) solution in chloroform, 14 μL of 1 mg/mL Oregon Green 488 1,2-dihexadecanoyl-sn-glycero-3-phosphoethanolamine (OG-DHPE, Invitrogen, Cat. No. 10154432) solution in chloroform was introduced. The mixture was then dried with a gentle flow of nitrogen to give a thin layer of lipids on the inner surface of a glass vial (Supelco, Cat. No. 27134). The vial was protected from light and kept under vacuum overnight at room temperature to remove the residual chloroform. Next, 1 mL of 18.2 MΩ cm water was added to the vial. The mixture was then vortexed for 1 min with a vortex mixer (PV-1, Grant-bio), after which the lipids were suspended in the water, forming multi-lamellar vesicles ("onions") presenting as a green-tinted cloudy suspension. The lipid suspension was sonicated for 20 min in an ice water bath with a 2 mm titanium probe (Sonicator microprobe 4423, Qsonica) mounted on a tip sonicator (Ultrasonic processor Q125, QSonica) with 60 % maximum power and 45/15 s on/off cycle, until it yielded a light-green clear suspension indicating the formation of small unilamellar lipid vesicles (SUVs) ~20-25 nm in size. The SUV suspension was centrifuged at

14,100g for 2 min (Centrifuge 5418 R, Eppendorf) to remove titanium particles, and the supernatant was retained and stored at 4 °C until use.

#### *Surface treatment of glass coverslips for the formation of supported lipid bilayers*

Glass coverslips (VWR, Cat. No. MENZBC026076AC40) were cleaned by sonication (Ultrasonic cleaner USC100T, VWR) for 5 min in 18.2 MΩ cm water, 5 min in 2-propanol (Sigma-Aldrich, Cat. No. 34863-2.5L), and 5 min in 18.2 MΩ cm water. The coverslips were then treated for 8 min with piranha solution, which is prepared by mixing 12 mL of 98% sulphuric acid (Merck, Cat. No. 5438270100) and 4 mL of 30% hydrogen peroxide solution (Sigma-Aldrich, Cat. No. 95321-100ML). Next, the piranha-treated coverslips were rinsed with excess 18.2 MΩ cm water and dried with a stream of nitrogen. A 50-well PDMS gasket (Merck, Cat. No. GBL103250) was brought into conformal contact with the coverslip. Another 50-well PDMS gasket was then aligned and brought into conformal contact with the top of the first gasket. The chambered coverslip was kept in a humidity chamber and used within 10 min. Specifically, the humidity chamber was made by adding 10 mL of 18.2 MΩ cm water to the bottom of an empty 250 μL pipette tip box (Rainin, Cat. No. 30389193).

#### *Formation of supported lipid bilayers*

The SUV suspension in water was mixed with phosphate-buffered saline (PBS, ThermoFisher, Cat. No. 10010023) at a ratio of 1:1 directly before use. The diluted SUV suspension (15 μL) was immediately added to each well of a chambered coverslip and incubated at 25 °C for 10 min in a humidity chamber. Next, 8 μL of the suspension was drawn from each well, followed by the addition of 8 μL of 18.2 MΩ cm water. This rinsing step was repeated three times. Then, 8 μL of the suspension was drawn from each well, after which 8 μL of PBS was added. This step was also repeated three times before the coverslip was ready for imaging.

#### *J8 LFA-1 Jurkat T-Cell sample preparation*

The J8 LFA-1 Jurkat T-cell line [1] used was kindly provided by the Davis lab in the University of Oxford. The cells were grown at 37 °C in 5% $CO_2$ in RPMI 1640 Medium, GlutaMAX™ Supplement, supplemented with 10% fetal calf serum, 10 mM HEPES, 1 mM sodium pyruvate, and 1% penicillin-streptomycin. The cell suspension was centrifuged at 2000 rpm for 2 min. The supernatant was removed, and the cell pallet was resuspended in 20 μL of 1x PBS. The cells were then stained with CellMask™ Orange Plasma membrane (ThermoFisher, Cat. No. C10045, 1:10000 in PBS) for 10 min at 37 °C, followed by the addition of 980 μL of 1x PBS. The suspension was resuspended in 1 mL of 1x PBS. The centrifugation-resuspension step was repeated three times to remove the excess dye. Finally, the cells were harvested and resuspended in 50 μL of 1x PBS, which was then added to the poly-L-lysine (PLL)-coated coverslip for imaging.

#### *Preparation of PLL-coated glass coverslip for CellMask™ Orange Plasma membrane stained-T cells imaging*

A 24 x 50 mm rectangular glass coverslip (VWR, Cat. No. 631-0146) was cleaned with argon plasma (PDC-002, Harrick Plasma) for 1 h. Two pieces of 8-well PDMS gasket (Merck, Cat. No. GBL103280) were aligned and attached to the coverslip. The glass surface was then treated with 1 mg/mL poly-L-lysine solution (Merck, Cat. No. P8920-100ML) for 5 min. Each well was washed with 1x PBS (0.02 μm-filtered) three times before the suspension of stained cells was added.

#### *Additional imaging conditions of CellMask™ Orange Plasma membrane stained-T cells*

The CellMask™ Orange Plasma membrane stained-T cells were imaged with different optical fibres installed on our home-built TIRF microscope with flat-field illumination setup. The images of the same FOV were acquired with three optical fibres (05806-1 Rev. A, CeramOptec; M42L01, Thorlabs; M122L02, Thorlabs) sequentially installed on the system. The sample was excited by the 561 nm laser. The acquired images were stacked and averaged.

#### *Preparation of oligonucleotides*

All oligonucleotides (Table S1) were purchased from ATDBio (Southampton, UK). They were synthesised on the 1.0 μmol scale and purified by HPLC, except for cy3B-labelled imaging strands, i.e. IS1 and IS2, which were purified by double HPLC. Lyophilised oligonucleotides were dissolved in 18.2 MΩ cm water (filtered by 0.02 μm filter (VWR, Cat. No. 516-1501)) to concentrations of 50-1000 μM as confirmed by $A_{260}$, aliquoted and stored at -20 °C.

| Code | Sequence (5' - 3') | Application |
|---|---|---|
| DBCO-DS2 | DBCO TEG-TTATCTACATATTTTTTTTTTTTTTTTTTTT | Labelling of mouse anti-rabbit antibody |
| IS2-cy3B | TATGTAGATC-cy3B | Cellular DNA-PAINT |
| Aptamer-DS1 | GCCTGTGGTGTTGGGGCGGGTGCGTTATACATCTA | AD-PAINT |
| IS1-cy3B | CTAGATGTAT-cy3B | AD-PAINT |

**Table S1. List of oligonucleotides and their applications. Underlined nucleotides are the complementary sequences that bind transiently to the docking strand and give a 9 bp DNA duplex during DNA-PAINT imaging. DBCO TEG – dibenzocyclooctyne tetraethylene glycol**

#### *Antibody labelling for DNA-PAINT imaging*

DBCO-DS2 was selectively conjugated on the carbohydrates of the $F_C$ region of the mouse anti-rabbit secondary antibody (Invitrogen, Cat. No. 31213, Lot No. UE2766762). To achieve this, the antibody was first functionalised using a SiteClick™ Antibody Azido Modification Kit (Invitrogen, Cat. No. S20026) according to the manufacturer's instructions. Briefly, 250 μg of antibody was concentrated to 3.4 mg/mL and buffer exchanged in the provided antibody preparation buffer. The antibody was then incubated overnight with β-galactosidase at 37 °C, followed by overnight coupling to UDP-GalNAz using β-1,4-galactosyltransferase (GalT) on the next day at 30 °C. The mixture was then purified by Amicon spin filter (50 kDa MWCO). The concentration of the azido-modified antibody was calculated by $A_{280}$ (1.9 mg/mL). With the azido-modified antibody, 10 molar equivalents of DBCO-DS2 was introduced for copper-free strain-promoted click reaction in 1×PBS. After overnight incubation at 37 °C, the excess oligonucleotide was removed using an Amicon spin filter (100 kDa MWCO) and the concentration of antibody and degree of labelling (3.2 docking strands per antibody) were determined by $A_{260}/A_{280}$. The purity and labelling efficiency were further confirmed using SDS-PAGE under reducing conditions (Figure S3).

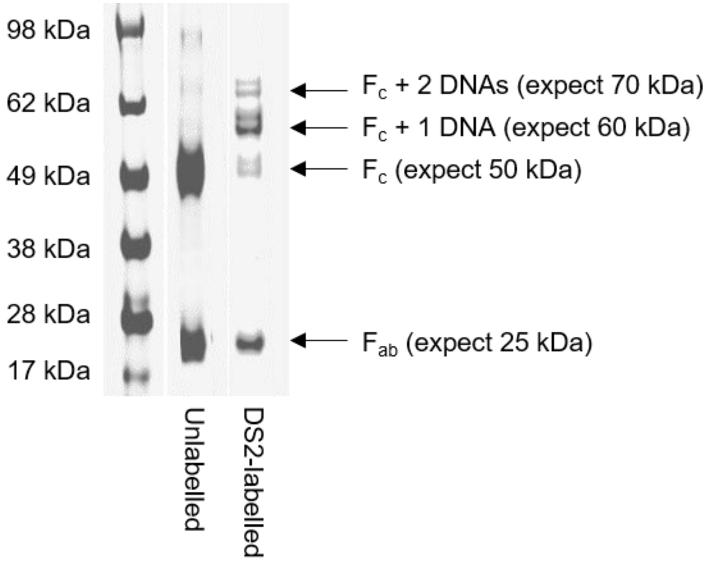


Figure S3. Characterisation of DNA-labelled mouse anti-rabbit antibody by SDS-PAGE under reducing conditions.

#### *HEK293T Cell sample preparation*

HEK293T cells were maintained in complete DMEM supplemented with 10% FCS, 100 U/mL penicillin, 100 µg/mL streptomycin and grown at 37°C and 5% $CO_2$. The cells were plated in µ-slide eight-well glass-bottom chambers (ibidi, Cat. No. 80827) pre-treated with 0.1 % poly-L-lysine (Merck, Cat. No. P8920) and allowed to adhere overnight in complete medium. The next day each well was incubated for ~30 seconds with 200 µL of extraction buffer containing 0.25% Triton and 0.1% glutaraldehyde in PEM buffer (80 mM PIPES, 5 mM EGTA, 2 mM $MgCl_2$, pH 6.8) at 37°C and the cells were subsequently fixed using 200 µL of fixation buffer containing 0.25% Triton and 0.5% glutaraldehyde in PEM buffer for 10 min at 37°C. The samples were then washed three times with PBS (Merck, Cat. No. 806544) and incubated with 0.1% $NaBH_4$ in PBS for 7 min at room temperature. This was followed by a 30-min blocking step using blocking buffer containing 0.5 % fish gelatin for dSTORM samples, with an addition of 1 mg/mL salmon sperm DNA (ThermoFisher, Cat. No. AM9680) for DNA-PAINT samples. The microtubules were immunostained overnight using a rabbit anti-tubulin antibody (abcam, Cat. No. ab18251, Lot No. GR3240348-1) diluted 1:300 in blocking buffer. The next day the cells were rinsed three times with PBS and subsequently incubated with either an Alexa-Fluor-647-labeled goat anti-rabbit IgG (H+L) (Invitrogen, Cat. No. A32733, Lot No. UH283999) at a concentration of 4 µg/mL in the blocking buffer for dSTORM, or a DS2-labelled mouse anti-rabbit antibody at a concentration of 2 µg/mL for DNA-PAINT, for 30 min at room temperature. Finally, samples were rinsed three times with 1xPBS before the introduction of the imaging solution.

#### *Imaging solution for dSTORM*

The imaging solution for dSTORM was prepared as reported [2]. Briefly, the following stock solutions were prepared.

- 0.1 M of tris supplemented with 20 mM NaCl, pH 8, filtered by 0.02 µm filter (VWR, Cat. No. 516-1501), stored at 4 °C (2x imaging solution of dSTROM)

- 25% glucose, stored at 4 °C (2.5x imaging solution of dSTORM)
- 1 M of cysteamine (Merck, Cat. No. 30070) in 0.36 M HCl, stored at 4 °C for no more than one week (20x imaging solution of dSTORM)
- GOD buffer (24 mM PIPES, 4 mM $MgCl_2$, 2 mM EGTA) at pH 6.8 and filtered by 0.02 µm filter, stored at 4 °C
- 20 mg/mL glucose oxidase from *Aspergillus niger* (Merck, Cat. No. G2133) in GOD buffer, centrifuge filtered with 0.22 µm filter (Merck, Cat. No. UFC30GV0S), flash frozen in liquid nitrogen, stored in -80 °C (40x imaging solution of dSTORM)
- 5 mg/mL catalase (Merck, Cat. No. C40) in GOD buffer, centrifuge filtered with 0.22 µm filter (Merck, Cat. No. UFC30GV0S), flash-frozen in liquid nitrogen, stored in -80 °C (125x imaging solution of dSTORM)

The final working imaging solution for dSTORM contains 0.5 mg/mL glucose oxidase, 40 µg/mL catalase, 50 mM cysteamine and 10% glucose in 50 mM of Tris supplemented with 10 mM NaCl at pH 8. This solution was prepared freshly and immediately before imaging.

#### *Preparation of recombinant α-Synuclein aggregates*

Wild type α-Synuclein was expressed, purified in *E. coli* and stored at -80 °C as described previously [3]. To remove pre-aggregation seeds, the solution was centrifuged at 91000*g* at 4 °C for 1 h by an ultracentrifuge (Optima TLX Ultracentrifuge, Beckman). The concentration of the supernatant was then determined by $A_{280}$ ($\varepsilon_{280}$ = 5960 $M^{-1}$ $cm^{-1}$). The supernatant was then diluted to 70 µM in 1xPBS supplemented with 0.01% $NaN_3$ (Merck, Cat. No. 71290) and incubated at 37 °C with shaking at 200 rpm for two months.

#### *Annealing of aptamer-DS1*

The stock solution of aptamer-DS1 (1000 µM) was diluted ten-fold into lithium cacodylate buffer, which contains 1 M of KCl (Breckland, EC No. 231-211-8, Stock Code: 0001276), 0.1 M of cacodylic acid (Merck, Cat. No. C0125), 0.1 M of lithium hydroxide (Merck, Cat. No. 909025) in 0.02 µm-filtered 18.2 MΩ cm water, pH 7.3. The aptamer-DS1 solution at 100 µM was heated to 95 °C for 10 min, and then cooled down slowly overnight to room temperature.

#### *AD-PAINT*

AD-PAINT was performed as we described previously [4]. Briefly, a 50 mm-diameter round coverslip (VWR, Cat. No. 631-0178) was cleaned with argon plasma (PDC-002, Harrick Plasma) for 1 h. A 50-well PDMS gasket (Merck, Cat. No. GBL103250) was cut into halves and then affixed to the coverslip. The slide was then treated with 0.02 µm-filtered (VWR, Cat. No. 516-1501) 1% Tween (Fisher Scientific, Cat. No. BP337-100, Lot No. 179118)/PBS (ThermoFisher, Cat. No. 10010023) for 1 h. It was then rinsed three times with 1xPBS (0.02 µm-filtered). Two-month alpha-synuclein (280 nM) was then introduced onto the slide and incubated for 5 min. After incubation at room temperature, the sample was removed, and the wells were filled with the imaging solution. To avoid evaporation over prolonged imaging, another clean coverslip was layered on top of the PDMS gasket.

#### *Imaging solution for DNA-PAINT*

For cellular DNA-PAINT imaging:

The imaging strand, IS2-cy3B, was in an oxygen-scavenging system PO+C. The oxygen-scavenging system was prepared as previously reported [5]. Briefly, the following stock solutions were prepared.

- PO+C buffer (10 mM Tris pH 7.5, 50 mM KCl and 20% glycerol), stored at 4 °C

- 38 mg/mL of pyranose oxidase from *Coriolus* sp. (Merck, Cat. No. P4234) in PO+C buffer (100x imaging solution for cellular DNA-PAINT imaging), centrifuge filtered with 0.22 µm filter (Merck, Cat. No. UFC30GV0S), flash-frozen in liquid nitrogen, stored in -80 °C
- 2 mg/mL of catalase (Merck, Cat. No. C40) in PO+C buffer, centrifuge filtered with 0.22 µm filter (Merck, Cat. No. UFC30GV0S), flash-frozen in liquid nitrogen, stored in -80 °C (100x imaging solution of cellular DNA-PAINT imaging)
- 100 mg of Trolox (Merck, Cat. No. 238813) was dissolved in 430 µL methanol, 345 µL NaOH (1 M) and 3.2 mL of 0.02 µm-filtered (VWR, Cat. No. 516-1501) 18.2 MΩ cm water, stored at -20 °C (100x imaging solution of cellular DNA-PAINT imaging)

The final working imaging solution for cellular DNA-PAINT imaging contains 5 nM of IS2-cy3B, 3.8 mg/mL pyranose oxidase, 20 µg/mL catalase and 0.25 mg/mL Trolox.

For AD-PAINT imaging:

The imaging solution was prepared as we previously reported [4]. Briefly, ultrapure grade thioflavin-T (Anaspec, Cat. No. AS-88306) was dissolved in 18.2 MΩ cm water at 100 µM as confirmed by $A_{412}$ ($\varepsilon_{412}$ = 31600 $M^{-1}cm^{-1}$). The thioflavin-T solution was then filtered by a 0.02 µm filter (VWR, Cat. No. 516-1501), stored at 4 °C in dark for no more than a month. The final working imaging solution for AD-PAINT contains 1 nM of IS1-cy3B, 100 nM of Aptamer-DS1 and 5 µM of thioflavin-T solution in PBS (ThermoFisher, Cat. No. 10010023).

*Super-resolution image reconstruction and data analysis*

The positions of the transient bindings between the imaging and docking strands or photo-switching of the fluorophores, 'blinkings', were determined by either using

- the PeakFit plugin, a plugin of the GDSC Single Molecule Light Microscopy package (http://www.sussex.ac.uk/gdsc/intranet/microscopy/imagej/gdsc_plugins) for imageJ/Fiji [6] with a typical 'signal strength' threshold of 40 and a precision threshold of 20 nm at a magnification as 8; or
- the thunderSTORM plugin [7] for imageJ/Fiji [6] with magnification as 10, B-spline order and magnitude as 3 and 2.0 respectively and fitting radius and initial sigma as 3 and 1.6 pixels respectively.

To automate super resolution reconstruction for multiple images with the PeakFit plugin in image J/Fiji, a script working on the python 3.7 platform (https://github.com/Eric-Kobayashi/SR_toolkit) was applied.

The resolution was determined by plotting an FRC curve with the FIRE plugin [8] for imageJ/Fiji [6] for the image super reconstructed by thunderSTORM and determining the spatial frequency at which the curve drops below 1/7 [8].

### 2. Theoretical calculations

*Contribution of small, square-core MMF to TIRF imaging*

For a beam focused at the back focal aperture to effectively contribute to an evanescent field, the size of the beam $S_b$ has to be smaller than the size of the objective lens TIRF annulus $S_a$.

$S_a$ is given by $f_B(NA - n_s)$ [9] where $f_b$ is the back focal length of the objective lens given by $F_L/M$ where $F_L$ is the focal length of the standard tube lens, and $M$ is the magnification of the objective lens. $NA$ is the numerical aperture of the objective lens, and $n_s$ is the refractive index of the sample (i.e. PBS buffer). For the imaging system used in this work, $F_L$ is 200 mm, $M$ is 100 – hence $f_B$ is 2 – $NA$ is 1.49, and $n_s$ is 1.335. Therefore, $S_a$ is 310 µm.

$S_b$ is given by $S_c F_k / F_{cl}$ where $S_c$ is the size of the fiber core, $F_k$ is the focal length of the Koehler lens, and $F_{cl}$ is the focal length of the beam collimator. For the imaging system used in this work, $S_c$ is 70 µm, $F_k$ is 125 mm, and $F_{cl}$ is 40 mm. Therefore, $S_b$ is 262.5 µm.
